# From Deer-to-Deer: SARS-CoV-2 is efficiently transmitted and presents broad tissue tropism and replication sites in white-tailed deer

**DOI:** 10.1101/2021.12.14.472547

**Authors:** Mathias Martins, Paola M. Boggiatto, Alexandra Buckley, Eric D. Cassmann, Shollie Falkenberg, Leonardo C. Caserta, Maureen H.V. Fernandes, Carly Kanipe, Kelly Lager, Mitchell V. Palmer, Diego G. Diel

## Abstract

Severe acute respiratory syndrome coronavirus 2 (SARS-CoV-2), the causative agent of coronavirus disease 2019 (COVID-19) in humans, has a broad host range, and is able to infect domestic and wild animal species. Notably, white-tailed deer (WTD, *Odocoileus virginianus*), the most widely distributed cervid species in the Americas, were shown to be highly susceptible to SARS-CoV-2 in challenge studies and reported natural infection rates approaching 40% in free-ranging WTD in the U.S. Thus, understanding the infection and transmission dynamics of SARS-CoV-2 in WTD is critical to prevent future zoonotic transmission to humans and for implementation of effective disease control measures. Here, we demonstrated that following intranasal inoculation with SARS-CoV-2, WTD fawns shed infectious virus up to day 5 post-inoculation (pi), with high viral loads shed in nasal and oral secretions. This resulted in efficient deer-to-deer transmission on day 3 pi. Consistent a with lack of infectious SARS-CoV-2 shedding after day 5 pi, no transmission was observed to contact animals added on days 6 and 9 pi. We have also investigated the tropism and sites of SARS-CoV-2 replication in adult WTD. Infectious virus was recovered from respiratory-, lymphoid-, and central nervous system tissues, indicating broad tissue tropism and multiple sites of virus replication. The study provides important insights on the infection and transmission dynamics of SARS-CoV-2 in WTD, a wild animal species that is highly susceptible to infection and with the potential to become a reservoir for the virus in the field.

**Author summary:** The high susceptibility of white-tailed deer (WTD) to SARS-CoV-2, their ability to transmit the virus to other deer, and the recent findings suggesting widespread SARS-CoV-2 infection in wild WTD populations in the U.S. underscore the need for a better understanding of the infection and transmission dynamics of SARS-CoV-2 in this potential reservoir species. Here we investigated the transmission dynamics of SARS-CoV-2 over time and defined the major sites of virus replication during the acute phase of infection. Additionally, we assessed the evolution of the virus as it replicated and transmitted between animals. The work provides important information on the infection dynamics of SARS-CoV-2 in WTD, an animal species that - if confirmed as a new reservoir of infection - may provide many opportunities for exposure and potential zoonotic transmission of the virus back to humans.

## Introduction

The coronavirus disease 2019 (COVID-19) pandemic caused by the novel severe acute respiratory syndrome coronavirus 2 (SARS-CoV-2), has incurred over 260 million human cases and more than 5.2 million deaths around the world (https://covid19.who.int/). The newly identified SARS-CoV-2 is a positive sense, single stranded RNA virus that belongs to the *Sarbecovirus* subgenus, within the *Betacoronavirus* genus, of the family *Coronaviridae* [1]. Genome sequence analysis determined that SARS-CoV-2 is closely related to other sarbecoviruses identified in horseshoe bats in China [2–4]. Thus, this animal species is currently considered the most likely source of the ancestral virus from which SARS-CoV-2 originated [4,5]. However, given the genetic differences between SARS-like bat coronavirus and SARS-CoV-2, it has been proposed that another, as yet unidentified, animal species may have played a role as an intermediate host prior to zoonotic spillover of SARS-CoV-2 to humans [2,6].

Infection with SARS-CoV-2 involves binding of the viral spike (S) protein to its host receptor, the angiotensin-converting enzyme 2 (ACE2) [7,8]. Comparison of the human ACE2 protein demonstrated a high degree of homology between the human ACE2 protein and its orthologues in multiple animal species [9]. Among the species that the ACE2 protein shares a high degree of homology with human ACE2, are three species of deer, including Père David’s deer (*Elaphurus davidianus*), reindeer (*Rangifer tarandus*), and white-tailed deer - WTD (*Odocoileus virginianus*), suggesting potential susceptibility of these species to the virus [9]. Notably, a pioneer study by our group demonstrated that WTD are indeed highly susceptible to SARS-CoV-2 infection and shed high viral titers in the respiratory secretions [10]. The relevance of our findings demonstrating the susceptibility of WTD to SARS-CoV-2 has been recently highlighted by the detection of SARS-CoV-2 antibodies in wild WTD sampled in Michigan, Pennsylvania, Illinois, New York, and Texas which showed a seroprevalence of ~40% in the sampled populations [11,12]. Furthermore, two independent studies in Iowa [13] and Ohio [14] detected SARS-CoV-2 RNA in tissues and respiratory secretions collected from WTD, suggesting actual infections in free-ranging WTD. The high susceptibility of the species combined with the prevalence observed in free-ranging herds raise concerns about the possibility of WTD becoming a reservoir for SARS-CoV-2. Although current evidence suggests circulation of SARS-CoV-2 in wild WTD, several questions remain unanswered regarding the potential role of this species as a reservoir for the virus, including whether sustained transmission that would allow maintenance of the virus in the field occurs in free ranging WTD populations.

Here, we expanded on our original work demonstrating the susceptibility of WTD to SARS-CoV-2. We investigated the transmission dynamics of the virus between contact animals over time and assessed the viral evolution as it replicated and transmitted between WTD. Additionally, we dissected the patterns of virus shedding, tissue tropism, and identified the main sites of viral replication during infection.

## Results

### Subclinical SARS-CoV-2 infection in WTD results in high viral shedding in respiratory and oral secretions

We have recently shown that WTD are highly susceptible to SARS-CoV-2 infection [10]. In this study, we assessed replication, shedding patterns and transmission dynamics of SARS-CoV-2 in WTD. For this, fifteen 8-month-old fawns (n = 15) were maintained in a biosafety level 3 (Agriculture) (BSL-3Ag) facility at the National Animal Disease Center (NADC) and randomly allocated to six experimental groups (Fig 1A). Three fawns (n = 3) were maintained in room A as a control group (uninoculated), while six fawns (n = 6) were intranasally inoculated with SARS-CoV-2 (human isolate NYI67-20, 5 × 10^6.38^ TCID_50_.ml^−1^) and maintained in a separate room (room B or C) during the 22-day experimental period (Fig 1A). Six contact animals were allocated to three experimental groups that were maintained in pairs (n = 2) in separate rooms (rooms D, E and F). The contact and inoculated animals were moved into a clean room (room G or H) at three different time points post-inoculation (pi) (days 3 [n = 2], 6 [n = 2], and 9 [n = 2] pi), maintained in contact with the inoculated fawns for 48 h and were then moved back to their respective rooms (Fig 1A). All fawns in the study (controls and inoculated animals) were monitored daily for clinical signs and body temperature. No clinical signs were observed in any of the animals during the experimental period. A slight and transient increase in body temperature was noted in 5 out of 6 inoculated fawns on day 1 pi (Fig 2B), and in 1 out of 6 inoculated fawns on day 2 pi. Throughout the rest of the daily monitoring period (11 days pi) the body temperatures remained within normal physiological range (Fig 2B).

**Fig 1.**
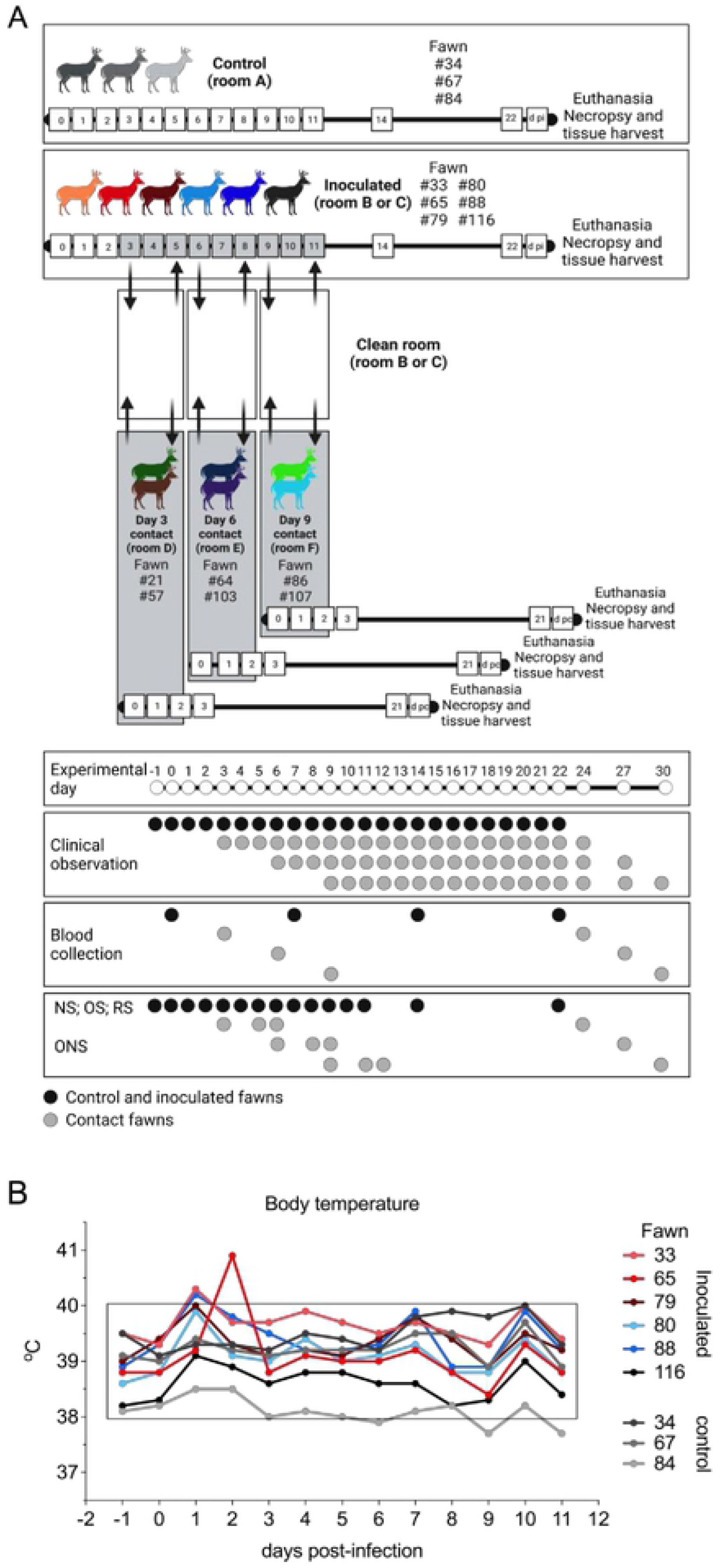
Infection and transmission study of SARS-CoV-2 in white-tailed deer. **A**) Schematic representation of the experimental design of infection/transmission study in WTD fawns. Fawns were kept in separate rooms in a biosafety level 3 (agriculture) (BSL-3Ag) facility. Three fawns were maintained as control (non-inoculated) in room A, and six fawns were inoculated intranasally with 5 × 10^6.38^ TCID_50_ of SARS-CoV-2 isolate NYI67-20 (lineage B1) and housed in room B or C for 22 days post-inoculation (d pi). To assess viral transmission dynamics, on day 3 pi two fawns (day 3 contact [pair 1]) were moved into a clean room (room C) where the inoculated fawns were also introduced. After 48 hours, day 3 contact fawns were transferred to a separate clean room (room D), where they were mainatined until euthanized and examined on 21 days post-contact (d pc). This process was performed with two additional fawns on day 6 pi (day 6 contact [pair 2]) and two fawns on day 9 pi (day 9 contact [pair 3]), which were housed in the separate rooms E and F, respectively. Respiratory nasal secretion, feces and serum were collected on the days indicated in the figure. **B**) Inoculated and control fawns were microchipped subcutaneously with temperature chips prior to the experiment, and monitored daily for body temperature starting on day 1 before inoculation until day 11 pi. Additionally, daily clinical observations were performed. Body temperatures are expressed in degrees Celsius. Rectangle highlights normal physiological body temperature range. NS = nasal swab; OS = oral swab; RS = rectal swab; ONS = oronasal swab.

**Fig 2.**
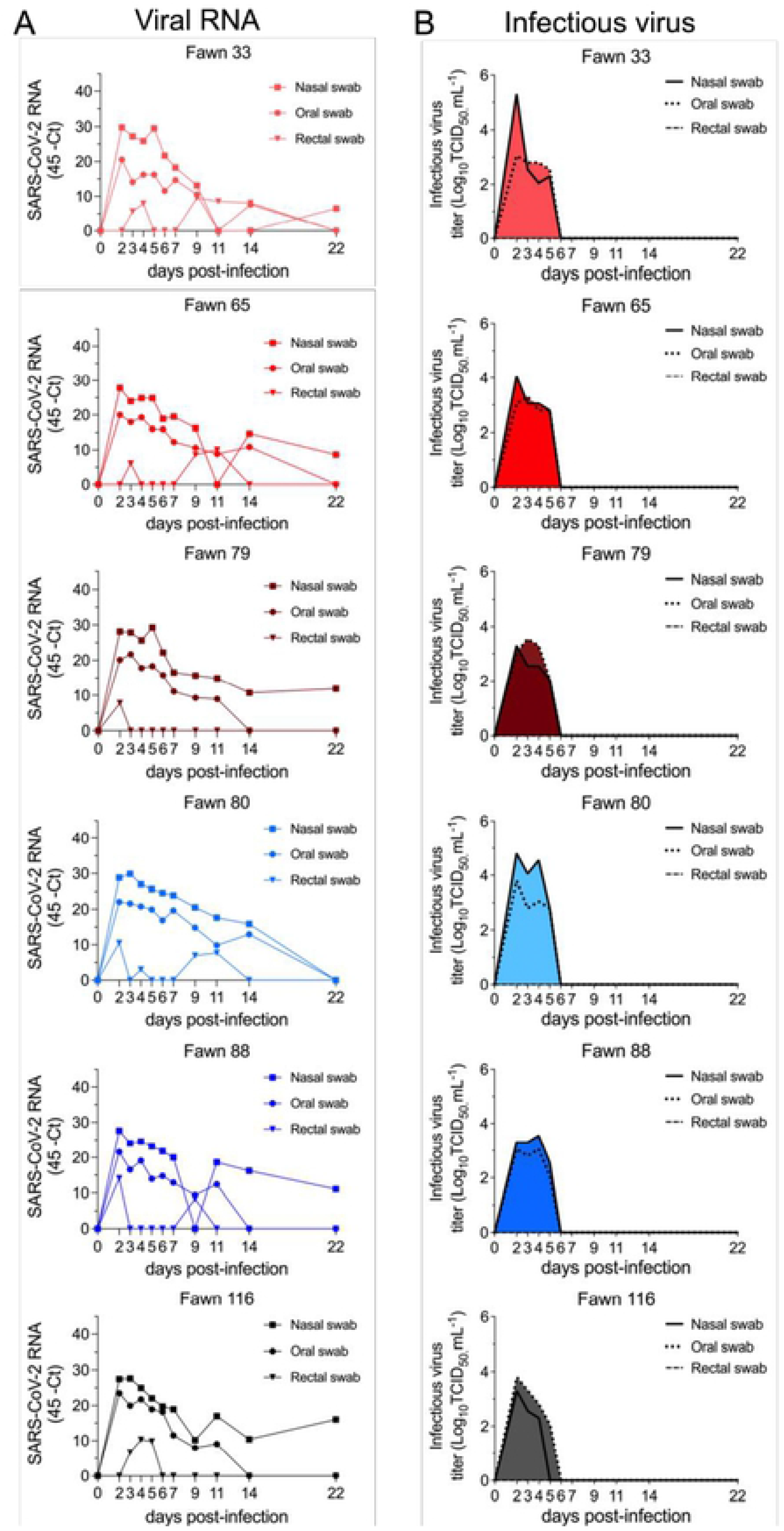
Viral shedding dynamics in nasal and oral secretions and feces following SARS-CoV-2 inoculation in white-tailed deer. **A**) SARS-CoV-2 RNA load assessed in nasal and oral secretions and feces of controls (animals no. 34, 67, 84, no shown) and six inoculated fawns (animals no. 33, 65, 79, 80, 88, and 116). Nasal, oral and rectal swabs collected on days 0, 2-7, 9, 11, 14, and 22 post-inoculation (pi) were tested for the presence of SARS-CoV-2 RNA by real-time reverse transcriptase PCR (rRT-PCR). **B**) Infectious SARS-CoV-2 load in nasal and oral secretions and feces was assessed by virus titration in RT-rPCR-positive samples. Virus titers were determined using end point dilutions and expressed as TCID_50_.ml^−1^. All the controls fawns (uninoculated) remained negative through the experimental period.

To monitor the infection and replication dynamics of SARS-CoV-2 in WTD, virus shedding was assessed following inoculation. Nasal and oral secretions and feces were collected using nasal (NS), oral (OS) and rectal swabs (RS) on days 0, 2, 3, 4, 5, 6, 7, 9, 11, 14 and 22 pi, from inoculated and control fawns. The samples were tested for the presence of SARS-CoV-2 by real-time reverse transcriptase PCR (rRT-PCR), and by virus isolation and titrations. All control fawns (uninoculated) remained negative through the experimental period. Viral RNA was detected between days 2 and 22 pi in nasal and oral secretions (Fig 2A) in the inoculated animals, with higher viral RNA loads detected between days 2 and 7 pi and decreasing thereafter through day 22 pi (Fig 2A). Shedding of viral RNA in feces were markedly lower compared to respiratory and oral secretions and was characterized by intermittent detection of low amounts of viral RNA (Fig 2A). The dynamics and duration of infectious SARS-CoV-2 shedding was also assessed in nasal and oral secretions and in feces. While no infectious virus was recovered from feces, high viral loads were detected in nasal and oral secretions (Fig 2B). All inoculated animals shed infectious SARS-CoV-2 between days 2-5 pi, with viral titers ranging from 2.0 to 5.3 and 2.0 to 3.8 log10 TCID_50_.ml^−1^, in nasal and oral samples, respectively. No infectious SARS-CoV-2 was recovered from samples collected after day 6 pi from any of the inoculated animals (Fig 2B).

### Efficient transmission of SARS-CoV-2 between WTD occurs shortly after infection

The transmission dynamics of SARS-CoV-2 between WTD was assessed by adding contact animals on days 3, 6 and 9 pi. SARS-CoV-2 transmission was only observed in fawns added on day 3 pi, as evidenced by detection of viral RNA and infectious virus in oronasal secretions from contact animals on days 2 and 3 pc (Fig 3A and B). Whereas contact animals added on days 6 and 9 pc did not become infected as evidence by the lack of viral RNA (Fig 3A and B). Notably, the two contact fawns added on day 3 pi shed infectious SARS-CoV-2 in oronasal samples on days 2 and 3 pc with titers ranging from 2.3 to 3.8 log10 TCID_50_.ml^−1^ (Fig 3B). Together these results indicate that SARS-CoV-2 productively infected and could be detected in the upper respiratory tract of contact fawns.

**Fig 3.**
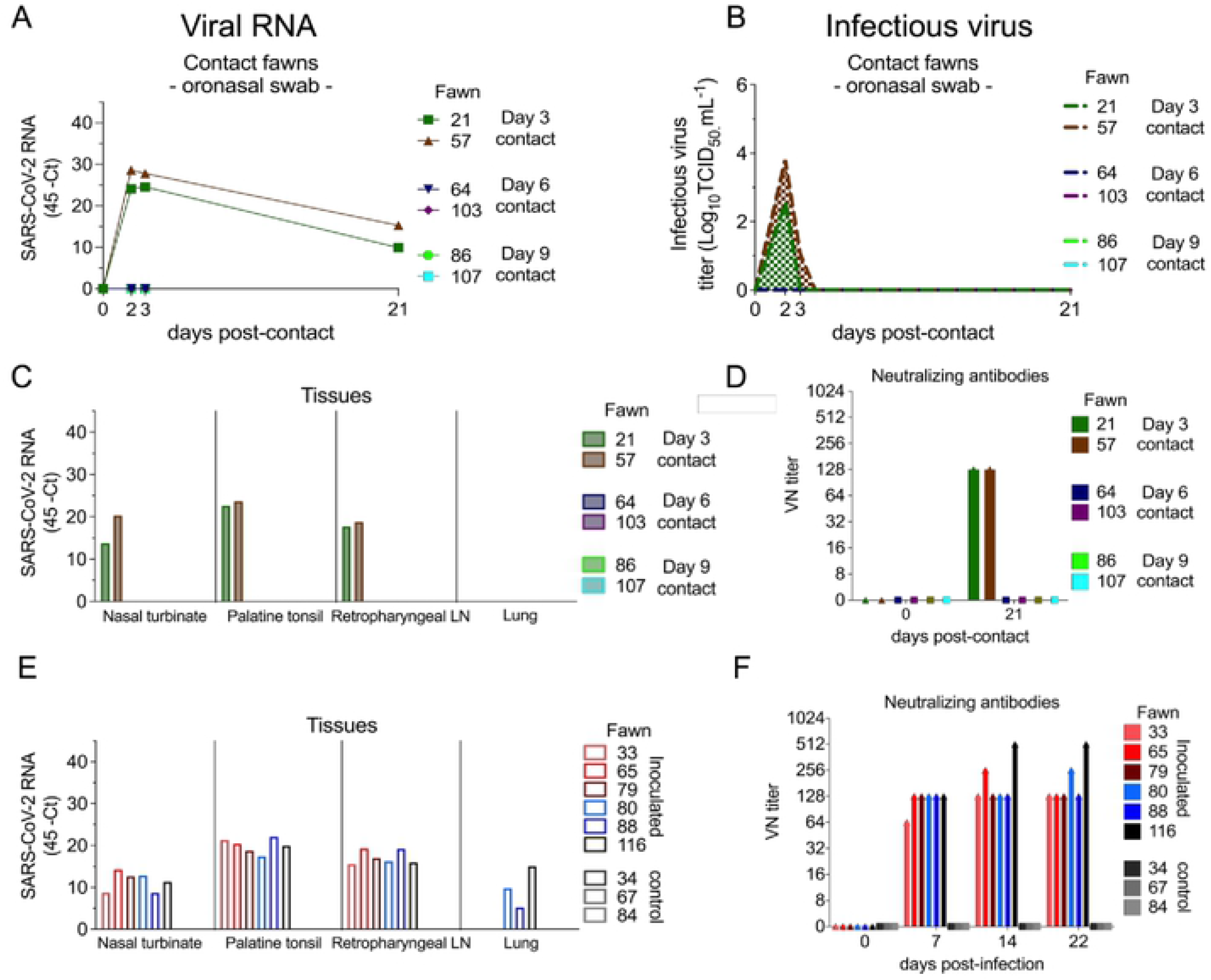
Viral load in oronasal secretions and feces, and seroconversion in contact white-tailed deer. **A**) Virus shedding (viral RNA load) assessed in oronasal secretions and feces of day 3-(animals no. 21 and 57), day 6-(animals no. 64 and 103), and day 9 contact (animals no. 86 and 107) fawns. Oronasal and rectal swabs collected on days 0 (before contact), days 2, 3 and 21 post-contact (pc) were tested for the presence of SARS-CoV-2 RNA by real-time reverse transcriptase PCR (rRT-PCR). **B**) Shedding of infectious SARS-CoV-2 in oronasal secretions and feces was assessed by virus titrations in rRT-PCR-positive samples. Virus titers were determined using end point dilutions and expressed as TCID_50_.ml^−1^. **C**) Tissue distribution of SARS-CoV-2 RNA assessed in nasal turbinate, palatine tonsil, retropharyngeal lymph node, and lung collected and processed for rRT-PCR 21 d pc of contact fawns. **D**) Antibody responses following contact of fawns with inoculated animals was assessed by virus neutralization (VN) assay. Neutralizing antibody titers were expressed as the reciprocal of the highest dilution of serum that completely inhibited SARS-CoV-2 infection/replication. **E**) Tissue distribution of SARS-CoV-2 RNA assessed in nasal turbinate, palatine tonsil, retropharyngeal lymph node, and lung collected and processed for rRT-PCR 22 days post-inoculation (d pi). **F**) Antibody responses following SARS-CoV-2 inoculation assessed by VN assay. Neutralizing antibody titers were expressed as the reciprocal of the highest dilution of serum that completely inhibited SARS-CoV-2 infection/replication.

Successful transmission and infection of SARS-CoV-2 to day 3 contact fawns was also confirmed by rRT-PCR performed in tissues including nasal turbinate, palatine tonsil, and medial retropharyngeal lymph nodes collected from these animals on day 21 pc (Fig 3C), which were positive for SARS-CoV-2 RNA. Importantly, viral RNA loads in these tissues were comparable to those observed in inoculated animals (Fig 3E). All tissues collected from contact animals added on days 6 and 9 of the experiment tested negative for SARS-CoV-2 by rRT-PCR (Fig 3C). Additionally, serological responses to SARS-CoV-2 (assessed by virus neutralization [VN] assay on day 21 pc) confirmed infection of day 3 contact animals. The two fawns added on day 3 seroconverted by day 21 pc (Fig 3D), presenting neutralizing titers comparable to those observed in inoculated animals (Fig 3F). Contact fawns added on days 6 and 9 pi, and all the controls animals, remained seronegative throughout the experiment.

### Low genetic diversity observed following SARS-CoV-2 replication and transmission in WTD

To investigate potential changes in the genome of SARS-CoV-2 following replication in WTD, genome sequence analysis was performed after infection and transmission to contact animals. SARS-CoV-2 genome sequences were obtained directly from nasal secretions from all the 6 inoculated fawns collected on days 3, 5, 7 and/or 9 pi and from oronasal secretions from the day 3 contact animals (n = 2) collected on days 2 and 3 pc. Whole genome sequence analyses of SARS-CoV-2 sequences recovered from all inoculated and contact fawns revealed no amino acid differences in the consensus SARS-CoV-2 genome in comparison to the genome sequence of the inoculum virus, a human SARS-CoV-2 isolate NYI67-20 of the B1 lineage (Fig 4A). Only minor variant viral populations with frequencies below 40% were observed (Fig 4B). Only four of these changes resulted in amino acid changes with one leading to truncation of ORF7 (Table S1).

**Fig 4.**
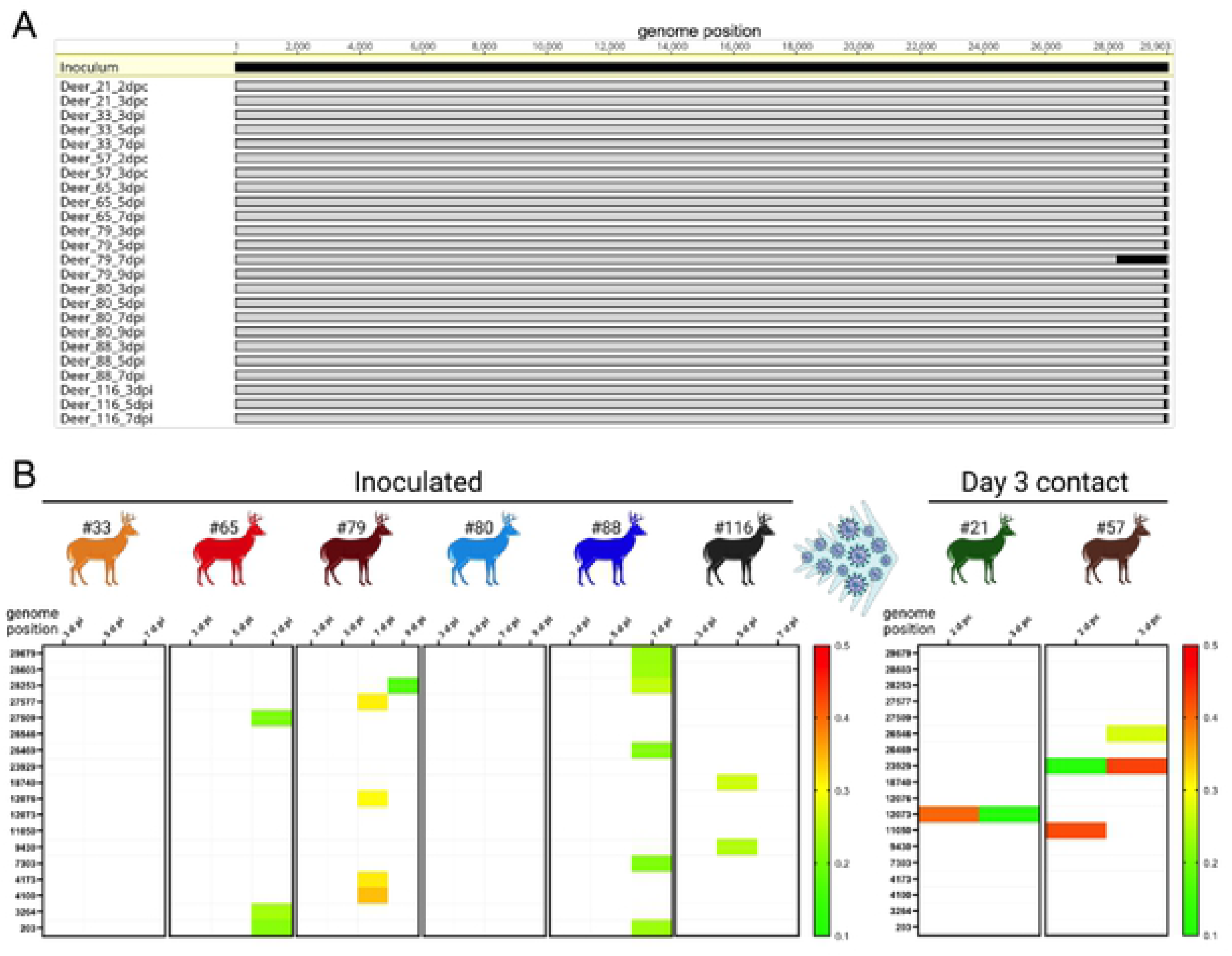
Low genetic diversity observed following SARS-CoV-2 replication and transmission in white-tailed deer. Genome sequences were obtained directly from nasal secretions from all the 6 inoculated fawns collected on days 3, 5, 7 and/or 9 post-inoculation (pi) and from oronasal secretions from the day 3 contact animals (n = 2) collected on days 2 and 3 post-contact (pc). **A**) Whole genome sequence analyses of SARS-CoV-2 sequences recovered from all inoculated and contact fawns revealed no amino acid differences in the consensus SARS-CoV-2 genome in comparison to the genome sequence of the inoculum virus isolate NYI67-20. **B**) Minor variant viral populations distributed throughout the virus genome from inoculated fawns on days 3, 5, 7 and/or 9 pi and from oronasal secretions from the day 3 contact animals (n = 2) collected on days 2 and 3 pc.

### Infection dynamics of SARS-CoV-2 in adult WTD is similar to that observed in WTD fawns

To gain insights into the sites of SARS-CoV-2 replication during acute infection, we conducted a second study in adult WTD. For this, eight deer (3-4 years of age; n = 8) were used and kept in the BSL-3Ag facility at the NADC. Two animals were maintained as controls (uninoculated) in room A, while six animals were inoculated intranasally with SARS-CoV-2 (human isolate NYI67-20, 5 × 10^6.38^ TCID_50_.ml^−1^ – same virus stock used in the transmission dynamics study) and housed in a separate room (room B) (Fig 5A). Following inoculation, all animals were monitored daily for clinical signs and body temperature was recorded daily until the day of euthanasia of each animal. No clinical signs were observed in any of the animals during the experimental period. Similar to inoculated fawns, a slight and transient increase in body temperature was noted in 4 of 6 inoculated deer on day 1 pi (Fig 5B). To assess viral tissue distribution and sites of virus replication, two inoculated deer were euthanized on day 2 and two on day 5 pi. The other two inoculated animals and two controls were euthanized on day 20 pi (Fig 5A). Viral RNA was detected until day 20 pi, or until the day of euthanasia (day 2 or 5 pi) in nasal secretions, and only sporadically in feces (Fig 5C-E). Similar to results obtained in fawns inoculated, high viral loads were detected in nasal secretions with titers ranging from 1.8 to 6.3 log10 TCID_50_.ml^−1^ in the early phase of infection (day 2 to 6 pi), while no infectious virus was detected in feces (Fig 5F-H). These results demonstrate similar clinical outcome, infection dynamics and shedding patters of SARS-CoV-2 in fawns (Fig 1B and 2A and B) and adult deer (Fig 5B-H).

**Fig 5.**
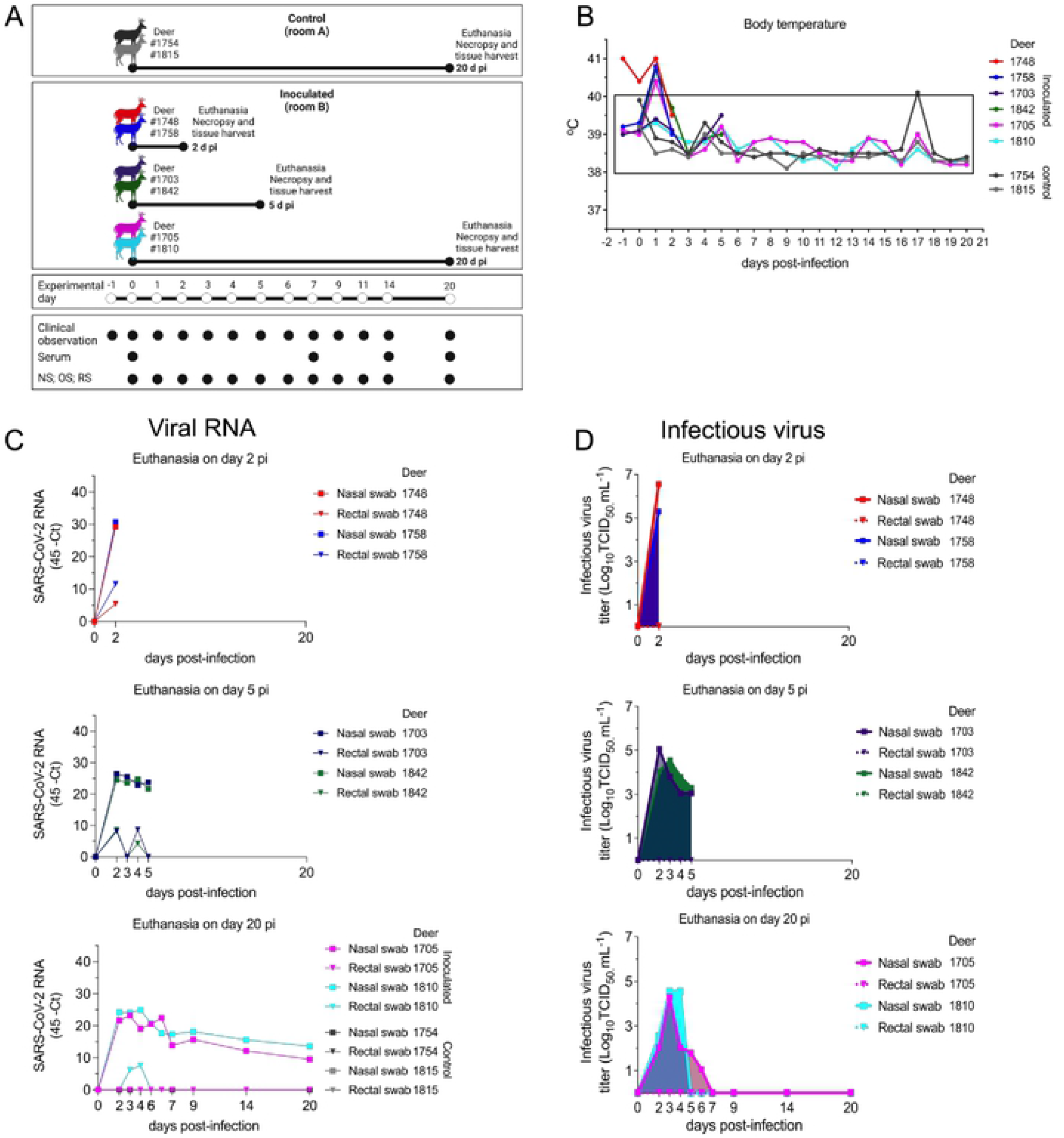
Pathogenesis and tissue tropism of SARS-CoV-2 in white-tailed deer. **A**) Schematic representation of the pathogenesis study design. Adult deer were kept in two rooms of a biosafety level 3 (agriculture) (BSL-3Ag) facility. Two deer were maintained as control (uninoculated) in a separate rom (room A), and six deer were inoculated intranasally with 5 × 10^6.38^ TCID_50_ of SARS-CoV-2 isolate NYI67-20 (lineage B1) and housed in room B. To assess viral tissue tropism, two deer were euthanized on day 2 post-inoculation (pi) (animals no.1748 and 1758), two on day 5 pi (animals no. 1703 and 1842), and two inoculated (animals no. 1705 and 1810) and two control (animals no. 1754 and 1815) deer were euthanized on day 20 pi. **B**) Inoculated and control deer were microchipped subcutaneously for temperature monitoring. Temperature and clinical signs were monitored daily starting on day 1 before inoculation until day 20 pi or until euthanasia. Body temperatures are expressed in degrees Celsius. **C-E**) SARS-CoV-2 RNA load assessed in respiratory secretions and feces by real-time reverse transcriptase PCR (rRT-PCR) in deer until days 2 (**C**), 5 (**D**), or 20 pi (**E**). **F-H**) Infectious SARS-CoV-2 assessed in respiratory secretions and feces assessed by virus titration in rRT-PCR-positive samples until days 2 (**F**), 5 (**G**), or 20 pi (**H**). Virus titers were determined using end point dilutions and expressed as TCID_50_.ml^−1^. All control deer (uninoculated) remained negative throughout the experimental period.

### SARS-CoV-2 presents a broad tissue tropism and replication sites during acute infection in WTD

To assess SARS-CoV-2 tissue tropism and replication sites, twenty-four tissues were collected and processed for rRT-PCR and infectious virus quantification. SARS-CoV-2 RNA was consistently detected in 17 out of 24 tissues sampled from both deer euthanized on day 2 pi, with the highest viral RNA loads detected in the nasal turbinate and palatine tonsil (Fig 6A). Similarly, from the tissues collected on day 5 pi, several tested positive by rRT-PCR, confirming a broad tissue distribution of SARS-CoV-2 RNA. As observed on day 2 pi, nasal turbinate and palatine tonsil were the tissues in which the highest viral RNA loads were detected on day 5 pi (Fig 6B). By day 20 pi, viral RNA loads decreased when compared to early times pi, and viral RNA was more consistently detected in lymphoid tissues such as palatine and pharyngeal tonsil, as well as medial retropharyngeal- and mandibular lymph nodes (Fig 6C). All tissues from the controls animals (uninoculated) tested negative by rRT-PCR (Fig 6C).

**Fig 6.**
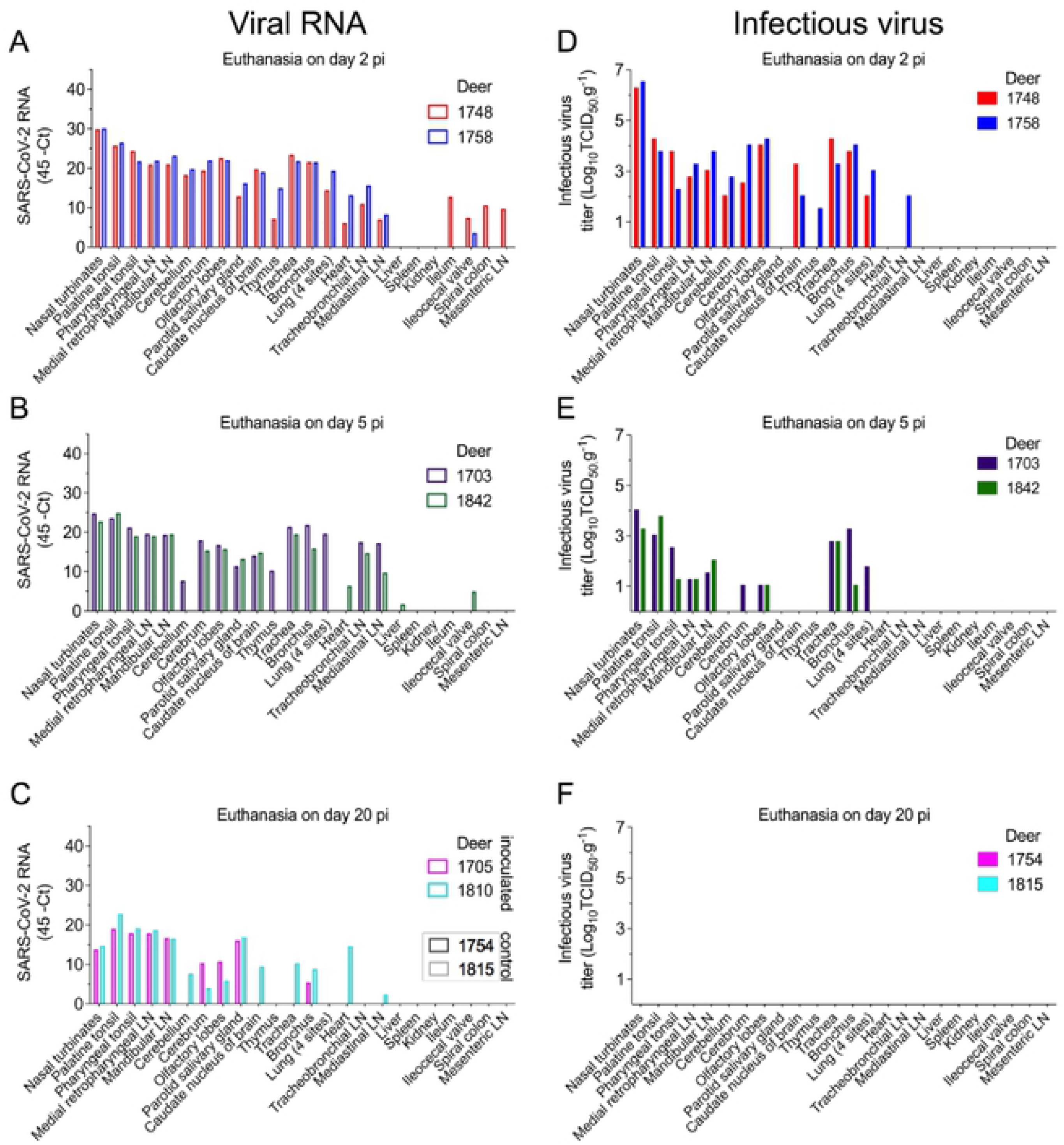
Tissue distribution of SARS-CoV-2 RNA and infectious virus in white-tailed deer. Tissues were collected and processed for real-time reverse transcriptase PCR (rRT-PCR) and virus titrations. **A**-**C**) Tissue distribution of SARS-CoV-2 RNA assessed in twenty-four tissues collected and processed for rRT-PCR in deer on days 2 (**A**), 5 (**B**), or 20 post-inoculation (pi) (**C**). **D**-**F**) Infectious SARS-CoV-2 in tissues assessed by virus titrations in RT-rPCR-positive samples obtained on days 2 (**D**), 5 (**E**) or 20 pi (**F**). Virus titers were determined using end point dilutions and expressed as TCID_50_.ml^−1^. All the controls fawns (uninoculated) remained negative through the experimental period.

Replication of SARS-CoV-2 was investigated in rRT-PCR positive tissues by viral titrations. Notably, infectious virus was consistently detected in a broad range of tissues on days 2 and 5 pi including nasal turbinate, tonsil (palatine and pharyngeal), lymph nodes (medial retropharyngeal, mandibular and tracheobronchial) and/or in the low respiratory tract tissues (trachea, bronchus and lung). Interestingly, infectious virus was also detected in olfactory lobe, caudate nucleus of brain, cerebrum, and cerebellum in both deer euthanized on day 2 pi (Fig 6D). On day 2 pi, viral titers ranged from 1.5 to 6.5 log10 TCID_50_.g^−1^ (Fig 6D), with the highest viral loads detected in the nasal turbinate of the two animals euthanized on day 2 pi (6.3 and 6.5 log10 TCID_50_.g^−1^) (Fig 6D). On day 5 pi, infectious virus was detected in 8 out of 24 tissues in both euthanized animals, with viral titers ranging between 1.0 to 4.0 log10 TCID_50_.g^−1^ (Fig 6E). Infectious virus was still detected in nasal turbinate (titers of 3.3 and 4.0 log10 TCID_50_.g^−1^) and in the olfactory lobes (titers 1.0 log10 TCID_50_.g^−1^) in the two deer euthanized on day 5 pi (Fig 6E). No infectious virus was detected in tissues collected from inoculated animals euthanized on day 20 pi (Fig 6F).

The replication of SARS-CoV-2 in tissues was confirmed by *in situ* hybridization (ISH) and immunohistochemistry (IHC) staining. Nasal turbinate, palatine tonsil, medial retropharyngeal lymph nodes, and lung from inoculated deer euthanized on days 2 and 5 pi were examined by ISH using the RNAscope ZZ technology and positive and negative sense RNA probes to detect SARS-CoV-2 genomic and subgenomic RNAs [10,15] and by IHC using a nucleoprotein (NP) specific mAb [15]. Moderate to abundant staining of viral RNA and NP was observed in nasal turbinate on days 2 and 5 pi; however, viral staining was less intense on day 5 pi (Fig 7 and 8). Viral staining was most pronounced in cells within the mucosal epithelial layer (Fig 7 and 8). Epithelial cells with contact to turbinate lumens were most often virus positive. Some cells in the middle and basilar portions of the epithelium were also sporadically stained. Additionally, moderate punctate staining of cells within the submucosal interstitial stroma, as well as cells associated with submucosal glandular or vascular elements were also observed (Fig 8). Viral RNA and antigen distribution in nasal turbinate epithelial cells was consistent with detection of high levels of expression of the virus receptor ACE2 and the TMPRSS2 protease in these cells (Fig 9A-C).

**Fig 7.**
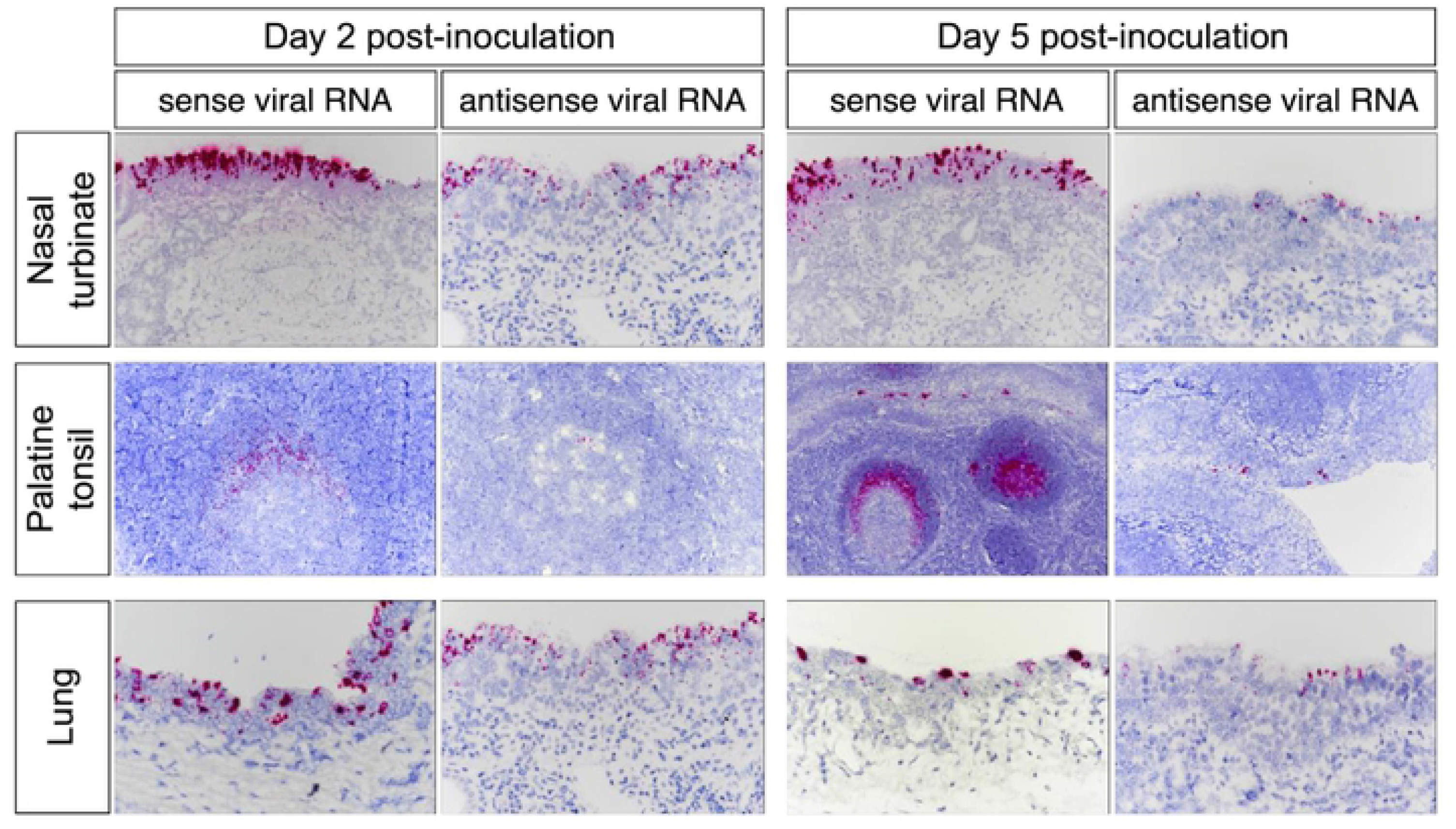
*In situ* hybridization (ISH) in tissues from white-tailed deer inoculated with SARS-CoV-2. Paraffin-embedded tissues were subjected to ISH using the RNAscope ZZ probe technology. Nasal turbinate, palatine tonsil and lung from deer on days 2 and 5 post-inoculation (pi). Intense labeling of viral RNA (all viral RNA) highlighted on the three tissues on days 2 and 5 pi. Labeling using the antisense genome probe demonstrate genome RNA replication in the nasal turbinate, palatine tonsil, and lung on day 2 pi and less intense on day 5 pi.

**Fig 8.**
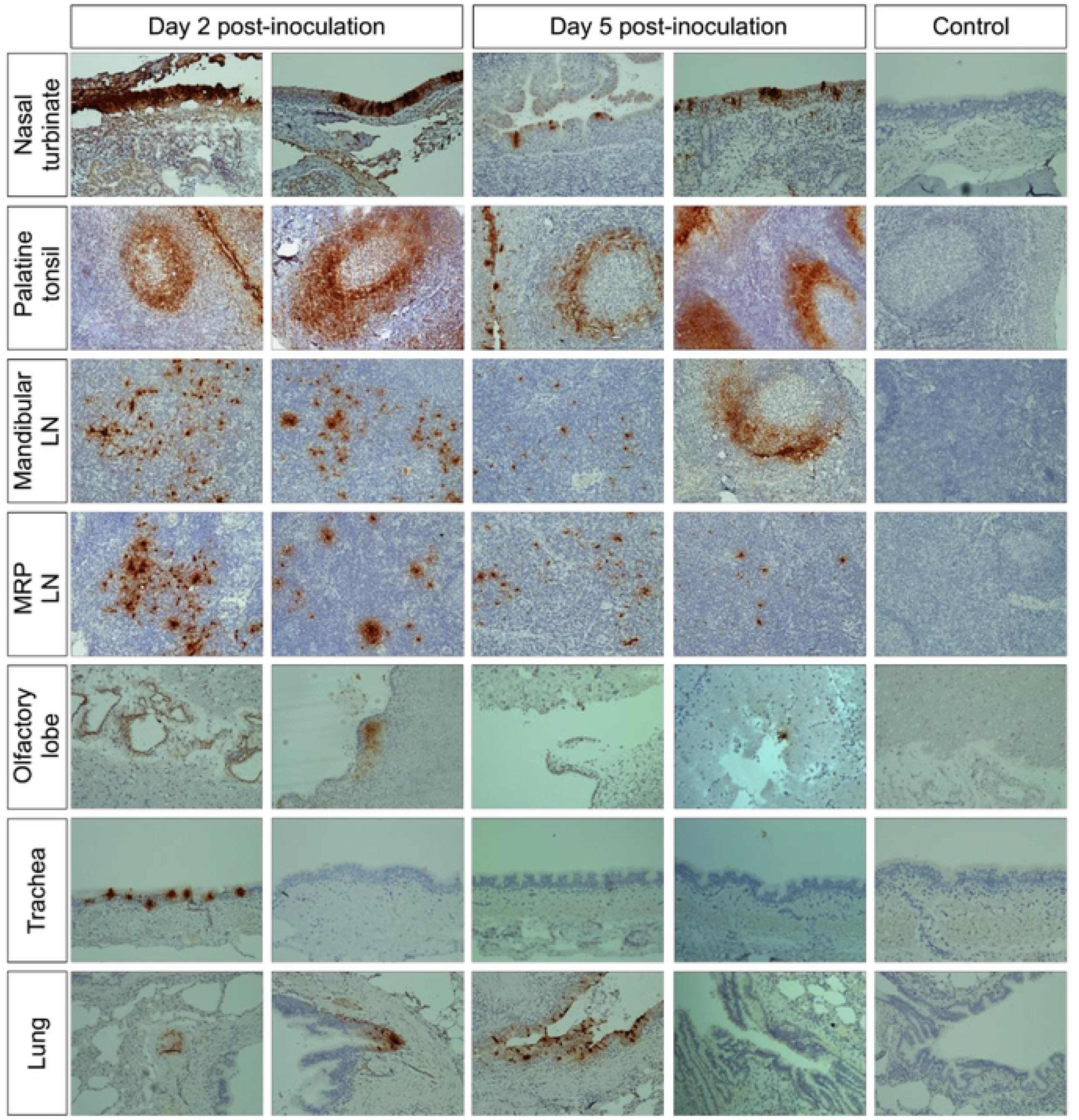
Immunohistochemistry (IHC) in tissues from white-tailed deer inoculated with SARS-CoV-2. SARS-CoV-2 labeling in the tissues of the inoculated WTD by IHC showing staining for the SARS-CoV-2 N protein (brown stain) in several tissues on days 2 and 5 post-inoculation (pi). Tissue sections were counter stained with hematoxylin. Tissues from a control (uninoculated) animal were included in all IHC stains.

**Fig 9.**
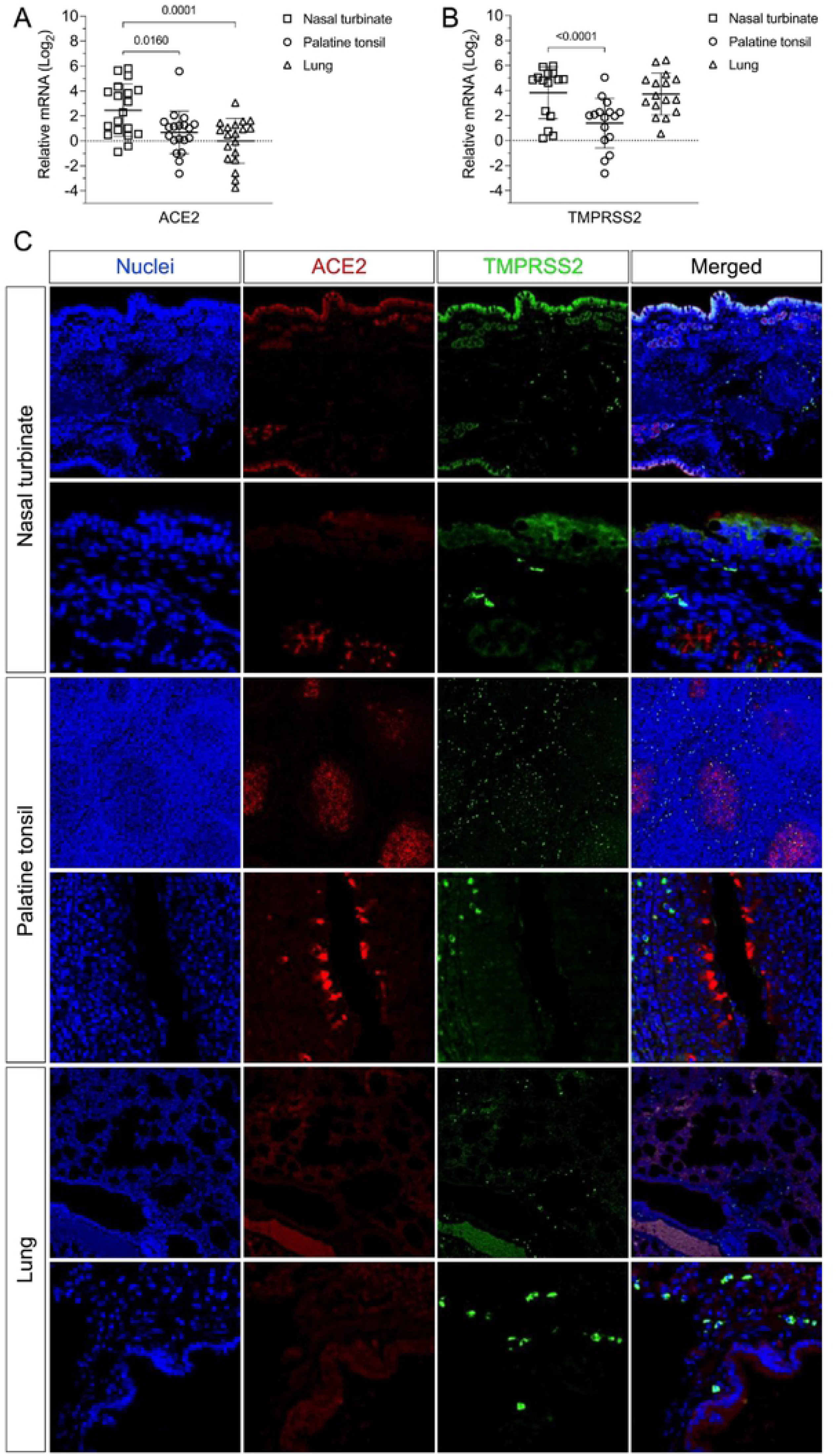
Expression of ACE2 and TMPRSS2 in the tissues of white-tailed deer. **A**-**B**) The levels of ACE2 and TMPRSS2 gene transcription in nasal turbinate, palatine tonsil and lungs were assessed by quantitative real-time reverse transcriptase PCR (qRT-PCR). Tissues from all 15 fawns from the transmission study (euthanized on days 21 or 22) and 4 deer from the pathogenesis study (euthanized on day 20) were included. Expression levels of ACE2 (**A**) and TMPRSS2 (**B**) in nasal turbinate, palatine tonsil and lung. **C)**Expression of ACE2 and TMPRSS2 proteins in the nasal turbinate, palatine tonsil, and lung are presented. Paraffin-embedded tissues from a control deer (uninoculated) were subjected to an immunofluorescence assay using a polyclonal antibody anti-ACE2 (red) and a monoclonal antibody anti-TMPRSS2 (green). Nuclear counterstain was performed with DAPI (blue).

Numerous tonsillar follicles and overlying epithelium presented virus specific RNA and NP staining on day 2 pi (Fig 7 and 8). Within the tonsillar crypt, virus positive epithelial cells were detected (Fig 7 and 8). Viral staining in lymphoid follicles was pronounced in the lymphocyte-rich cap and mantle regions of the tonsil, between tightly packed cells (Fig 7 and 8). On day 5 pi SARS-CoV-2 RNA- and NP staining in crypt epithelial cells, tonsillar reticular epithelium and lymphoid follicles was similar to that observed on day 2 pi. However, in addition to staining of the follicular cap and mantle regions, virus positive cells were also noted in portions of the central germinal centers (Fig 7 and 8). Notably, high levels of expression of ACE2 and TMPRSS2 were detected in crypt epithelial cells and in germinal centers in the tonsil (Fig 9C), two areas in which active virus replication was detected as demonstrated by antisense viral RNA staining (Fig 7 and 8). By day 20 pi, viral RNA staining was limited to germinal centers as was observed on days 2 and 5 pi (data not shown).

Viral staining in the mandibular and/or retropharyngeal lymph nodes was observed on days 2 and 5 pi, with the interfollicular and paracortical regions of these lymph nodes containing large foci of virus (Fig 8). Additionally, a lower number of positive cells with less intense staining was observed in the germinal center mantle and central regions (Fig 8). On day 20 pi, viral RNA staining within the medial retropharyngeal lymph node was characterized by low to moderate numbers of single positive cells within germinal centers (data not shown). Importantly, no evidence of viral replication (virus isolation) was observed in any of the tissues collected on day 20 pi (Fig 6F).

In the lung, moderate to abundant staining of bronchial epithelial cells was observed in both animals on day 2 pi (Fig 7 and 8) and in only one animal euthanized on day 5 pi. Large foci of virus positive cells (indicative of replication) were scattered throughout bronchial epithelium with infrequent staining of individual cells within the submucosa. Staining was absent in bronchioles, alveoli, or interstitial regions. Viral staining was also observed in tracheal epithelial cells of one animal on day 2 pi, with foci of intense virus staining observed in the ciliated epithelium (Fig 8). In general, virus-specific staining was more broadly distributed and more intense on day 2 pi with a noticeable decrease in the number of positive cells and staining intensity being observed on day 5 pi. Together these results demonstrate active SARS-CoV-2 replication in the upper and lower respiratory tract and lymphoid organs of WTD. Detection of higher viral loads in URT are consistent with higher expression levels of ACE2 and TMPRSS2 in nasal turbinate, when compared to tonsil and lung tissues (Fig 9A-C).

### Histological changes associated with SARS-CoV-2 replication

Tissue samples were processed for standard histological examination. Rhinitis characterized by submucosal lymphoplasmocytic infiltrates and frequent mucosal exudation of neutrophils (Fig 10) was observed in the nasal turbinate. In lymphoid organs (tonsil and lymph nodes), follicles exhibited moderate lymphoid depletion and lymphocytolysis (Fig 10). Tonsil also contained multifocal hemorrhages in crypt epithelium, crypts with variable numbers of neutrophils and cell debris, and congested vasculature filled with neutrophils. In the lungs, diffuse congestion of alveolar capillaries and multifocal hemorrhage in the submucosa of larger airways was observed (Fig 10). Multifocally, there were perivascular and interstitial lymphohistiocytic infiltrates. Occasionally, there were foci of mild increased alveolar macrophages and type II pneumocyte hyperplasia. In general, these histological changes were more pronounced in days 2 and 5 pi, with notable changes in histological features and extension observed by the end of the experiment on day 20 pi (data not shown).

**Fig 10.**
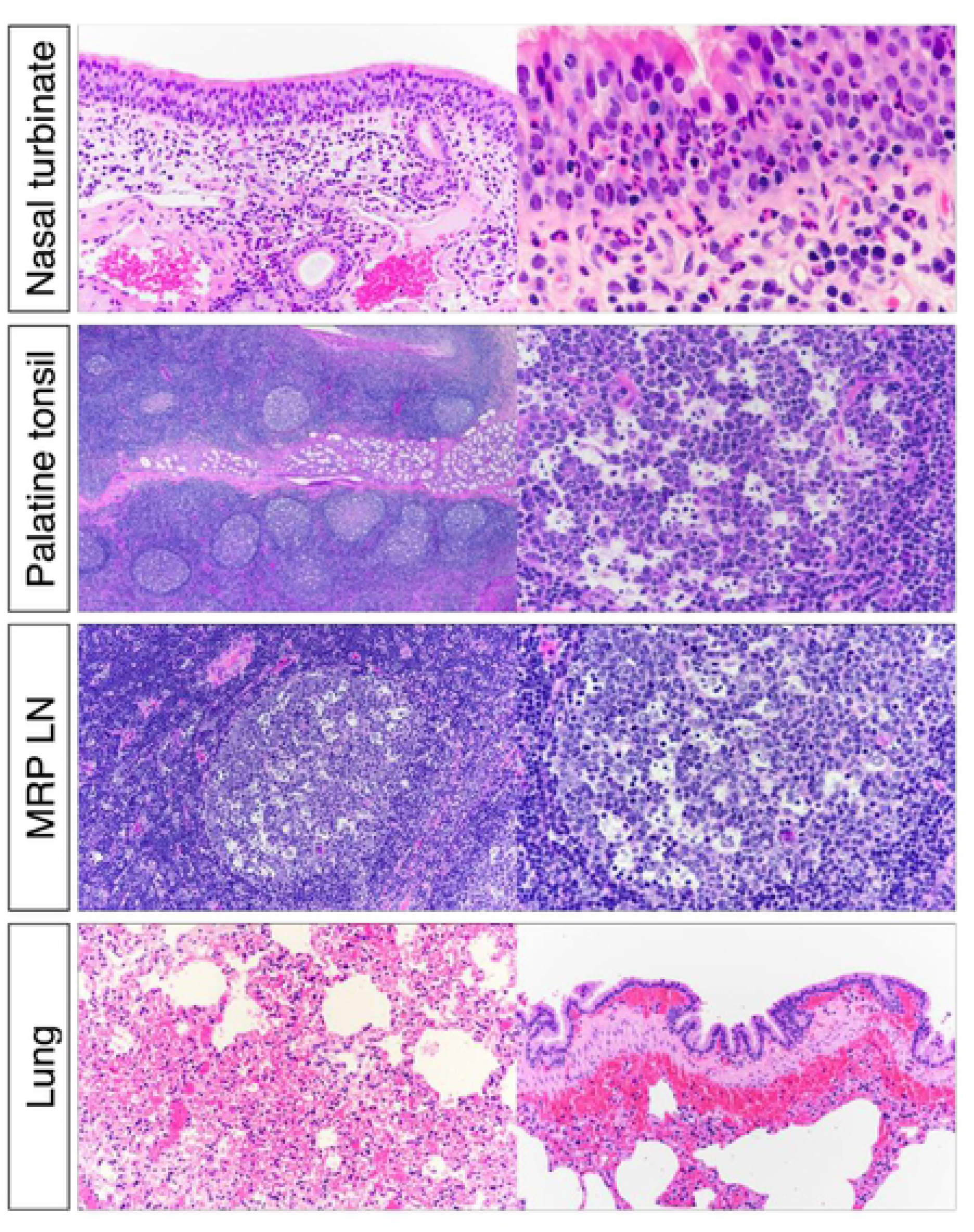
Histological examination of nasal turbinate from white-tailed deer inoculated with SARS-CoV-2. Upper respiratory tract (URT) after inoculation, there was submucosal lymphoplasmacytic infiltrate and frequent mucosal exudation of neutrophils in nasal turbinate. Numerous germinal centers of the palatine tonsils were characterized by lymphoid depletion and lymphocytolysis. Within the lung there was diffuse congestion with multifocal submucosal hemorrhages associated with larger bronchi.

## Discussion

Here, we characterized the window in which SARS-CoV-2 is transmitted through contact in WTD and defined important aspects of the infection dynamics and pathogenesis of the virus in this species, including its tissue tropism and sites of replication during acute infection.

The transmissibility of SARS-CoV-2 is a major factor that contributed to the establishment of the virus in the human population worldwide [16,17]. In WTD, we and others have shown efficient transmission of the virus to non-contact [10] or contact animals [18]. In this study, we investigated the patterns and dynamics of virus shedding and transmission to contact animals and demonstrated that infectious SARS-CoV-2 is shed mainly in respiratory and oral secretions during the first 5 days pi. Consistent with this, efficient transmission of the virus to contact animals only occurred during the period in which the animals were shedding infectious SARS-CoV-2, as evidenced by productive infection of day 3 contact animals, but not of animals added on days 6 and 9 pi. Interestingly, no infectious virus was recovered from feces collected from inoculated animals between days 2 to 22 pi. These observations are relevant when considered in the context of recent studies demonstrating infection and circulation of SARS-CoV-2 in wild WTD in the U.S. [11–14]. Our findings suggest that transmission of SARS-CoV-2 in WTD is likely to occur mainly through respiratory or oral secretions, and the fact that no infectious virus was recovered from feces further suggests that efficient transmission of the virus would require close contact between infected and susceptible animals. Should WTD be confirmed a reservoir for SARS-CoV-2, the need for close contact and the short infectious period would limit potential opportunities for transmission of the virus to other susceptible wild animal species (e.g. raccoons, and deer mice [19–21]) and to humans.

To better understand host selective pressures on the SARS-CoV-2 genome, we analyzed whole genome sequences obtained from inoculated and contact WTD following replication and transmission of the virus over the course of the 22-day experiment. Notably, no amino acid changes were observed in the SARS-CoV-2 consensus genome throughout the experimental period, suggesting low host-selective pressure(s) on the virus in WTD. Similar observations were reported in SARS-CoV-2 sequences obtained from free ranging WTD [13,14], which revealed the circulation of closely related viruses, with all sequences recovered from WTD being identical to sequences from SARS-CoV-2 obtained from humans in the same geographic region(s). This contrasts with what has been described in other animal species (e.g. tigers, lions, mink, cats, dogs, hamsters and ferrets) naturally and/or experimentally infected with the virus, in which characteristic mutations mainly in the S protein have been observed shortly after infection [22–25]. The findings in WTD can be partially explained by the high degree of homology of the human and deer ACE2 proteins [9] – the entry receptor for the virus – which may favor interaction of the S protein with the cellular receptor and result in lower selective pressure for virus entry and spread to target cells in secondary sites of replication. Importantly, when we analyzed the viral subpopulations recovered from WTD, we observed the presence of low frequency (<40%) variant viruses containing mutations at different loci dispersed throughout the genome. These variants, however, did not become established over the course of the experiment. The possibility, however, that beneficial viral mutations become fixed and variant strains emerge in this animal species, cannot be formally excluded.

Other important aspects of SARS-CoV-2 infection dynamics and pathogenesis that were investigated in our study are its tissue tropism and sites of replication during acute infection. Our results demonstrate a broad tissue tropism of the virus, with active replication taking place in respiratory (nasal turbinate, trachea, bronchus and lung), lymphoid (palatine and pharyngeal tonsil, retropharyngeal, mandibular and tracheobronchial lymph nodes and thymus) and central nervous tissues (olfactory lobes, cerebrum, caudate nucleus of brain and cerebellum). High viral loads were observed in nasal turbinates and lymphoid tissues associated with the oral cavity (tonsils, retropharyngeal and mandibular lymph nodes) or respiratory system (tracheobronchial lymph nodes) on days 2 and 5 pi. The high amounts of infectious virus recovered from nasal turbinate and tonsils (viral titers ranged from 3.3 to 6.5 log10 TCID_50_.g^−1^) are consistent with intense viral RNA and N protein staining observed *in situ* and with the viral load and shedding patterns detected in respiratory secretions. This indicates that nasal turbinate and tonsil are likely the primary sites of viral replication during infection in WTD, which is similar to observations in other animal species and in humans [19,26–29]. Following infection and replication in these sites, the virus likely disseminates to draining lymph nodes, the lower respiratory (trachea and lung) tract and the CNS. Efferent lymphatics from the palatine tonsil, for example, drain directly to the medial retrophayrngeal lymph nodes, which would explain the consistent involvement of that lymph node in the current study. Notably, higher viral loads and replication levels observed in nasal turbinate and tonsil, when compared to the levels observed in the lung for example, are consistent with distribution and higher expression levels of ACE2 and TMPRSS2 proteins in upper respiratory tissues.

Consistent with lack of virus shedding in feces, no viral RNA was detected in any of the gastrointestinal (GI) tissues collected in our study, indicating that the GI tract is not a major target of SARS-CoV-2 in WTD. An interesting finding of our study was the ability to isolate SARS-CoV-2 in the CNS of inoculated animals and may be suggestive of replicating virus in these tissues. A similar tropism has been described in transgenic hACE2 expressing mice [30–32], and deer mice [20,21], in which neural invasion has been attributed to the olfactory nerve. Importantly, neurological signs following SARS-CoV-2 have also been described in humans suggesting neuroinvasion [33–35], however, WTD in our study did not present clinical signs of neurological infection. Together our findings demonstrate that SARS-CoV-2 is able to reach and replicate in several tissues in WTD, remarkably in the nasal turbinate, tonsil and lymph nodes associated with the oral cavity and respiratory tract. The results presented here provide a detailed overview of the tissue tropism and replication sites of the virus in this highly susceptible animal species.

Our study extends the knowledge regarding SARS-CoV-2 infection dynamics in WTD, an animal species with the potential to serve as a reservoir for the virus in the field. We defined that SARS-CoV-2 has a short window of infectivity and transmissibility in WTD, which is similar to what has been described in humans and other animal species [19,21,27,29,36–38], and demonstrated that the virus presents a broad tissue tropism and multiple replication sites in this species. This information is critical to assess the relevance of natural SARS-CoV-2 infections in WTD populations and the potential risk of zoonotic spillback into humans. In our initial study demonstrating the susceptibility of WTD to SARS-CoV-2 [10], we suggested that deer and other cervids should be considered in investigations conducted to identify the origin and potential intermediate host species that may have served as the link host reservoir to humans. We had also highlighted that the high susceptibility of this species to SARS-CoV-2 combined with the constant interactions with humans could lead to opportunities for human-WTD transmission. This possibility has been recently confirmed by studies demonstrating natural SARS-CoV-2 infections in free-ranging WTD in the U.S. [11–14]. Several questions regarding these findings remain unanswered, perhaps the most important ones are: 1) Can WTD maintain the virus in the wild?; and 2) Can infected WTD transmit the virus back to humans? Our findings provide significant insights into the infection and transmission dynamics of SARS-CoV-2 in WTD. Such information is critical to design appropriate epidemiological studies to help answering important questions regarding the role and potential impact of WTD in the future of the COVID-19 pandemic, and most importantly to confirm or rule out the potential role of this species as a reservoir for the virus in the field.

## Material and Methods

### Cells and Virus

Vero E6 (ATCC^®^ CRL-1586™), and Vero E6/TMPRSS2 (JCRB Cell Bank, JCRB1819) were cultured in Dulbecco’s modified eagle medium (DMEM) supplemented with 10% fetal bovine serum (FBS), L-glutamine (2mM), penicillin (100 U.ml^−1^), streptomycin (100 μg.ml^−1^) and gentamycin (50 μg.ml^−1^) for both cell lines. While the culture media used to keep the Vero E6/TMPRSS2 was supplemented with geneticin (1 mg.ml^−1^). The cell cultures were maintained at 37 °C with 5% CO_2_. The human SARS-CoV-2 isolate NYI67-20 (lineage B1) was propagated in Vero E6 cells. Low passage virus stock (passage 3) were prepared, cleared by centrifugation (2000 × *g* for 15 min) and stored at −80 ^o^C. The whole genome sequence of the virus stock was determined to confirm that no mutations occurred during passages in cell culture. A viral suspension containing 10^6.38^ 50% tissue culture infectious dose per ml (TCID_50_.ml^−1^) was used for all *in vitro* and *in vivo* experiments.

### Animal studies

All animal work and procedures performed in this study were approved by the National Animal Disease Center (NADC) Institutional Animal Care and Use Committee for both the fawn study (protocol ARS-2020-902) and the adult deer study (protocol ARS-2020-861). White-tailed deer fawns (n = 15) with approximately 8 months-of-age were obtained from a captive deer facility in Hillman, Michigan, USA. Adult deer (3-4 years-old) seronegative to SARS-COV-2 were obtained from a breeding herd maintained at the NADC in Ames, IA. Animals were housed in a biosafety level 3 (Agriculture) (BSL-3Ag) facility at NADC, and allowed to acclimate for at least 2 weeks prior to the initiation of the study. All animals were microchipped (subcutaneously; SC) for identification and body temperature monitoring. All fawn and adult deer were sampled and screened for SARS-CoV-2 RNA by real-time reverse transcriptase PCR (rRT-PCR) in oronasal secretions and by virus neutralization assays prior to virus inoculation.

### Transmission dynamics of SARS-CoV-2 in white-tailed deer

After acclimation, six fawns were inoculated intranasally with 5 ml (2.5 ml per nostril) virus suspension containing 10^6.38^ TCID_50_.ml^−1^ of SARS-CoV-2 isolate NYI67-20 (lineage B1) and maintained in room B. Three fawns (control) were maintained as non-inoculated control animals in a separate room (room A). Body temperatures were recorded daily. Nasal swab (NS) and rectal swabs (RS) were collected on days 0, 2, 3, 4, 5, 6, 7, 9, 11, 14 and 21 post-inoculation (pi) (Fig 1A). Upon collection, flocked swabs were placed individually in sterile tubes containing 2 ml of viral transport media (MEM with 1,000 U.mL^−1^ of penicillin, 1,000 µg.mL^−1^ of streptomycin, and 2.5 µg.ml^−1^ of amphotericin B) and stored at −80 °C until analysis. Blood was collected through jugular venipuncture in serum separator tubes on days 0, 7, 14, and 22 pi. The tubes were centrifuged for 10 min at 1200 × *g* and serum was aliquoted and stored at −80 °C until analysis. To assess transmission, on day 3 pi, two fawns (contact pair 1) were bled, oronasal swab (ONS) collected, and introduced into a clean room (room C) along with the inoculated fawns. After the inoculated fawns were moved from room B, it was power washed, disinfected, and remained empty for 24 h until the next exposure. After 48 hours, contact pair 1 fawns were sampled again (ONS) and placed into a separate room until necropsy on 21 days post-contact (d pc). This contact process was repeated using cleaned room B or C with two fawns (contact pair 2), which were added on day 6 pi and two additional fawns (contact pair 3) added on day 9 pi (Fig 1A). For contact animals, swab samples were collected in viral transport media on 0, 2, 3, and 21 d pc. Blood was collected through jugular venipuncture using serum separator tubes on days 0 and 21 pc. Serum from the serum separator tubes was aliquoted and stored at −20 °C until analysis. Fawns were humanely euthanized on day 22 pi and 21 pc (inoculated and contact animals, respectively) with an intravenous dose of barbiturate (Fatal-Plus^®^ Solution, Vortech Pharmaceuticals, Dearborn, MI) according to the manufacturer label dose. Following necropsy, multiple tissues including: nasal turbinate, palatine tonsil, medial retropharyngeal lymph node, and lung [4 sites], were collected. Samples were processed for real-time reverse transcriptase PCR (rRT-PCR) and virus isolation (VI) were individually bagged, placed on dry ice, and transferred to a −80 °C freezer until testing.

### Tissue tropism of SARS-CoV-2 in white-tailed deer

After acclimation, six adult deer were inoculated intranasally with 5 ml (2.5 ml per nostril) of 10^6.38^ TCID_50_.ml^−1^ of SARS-CoV-2 isolate NYI67-20. Two animals were maintained as non-inoculated controls. Body temperatures were recorded daily. NS and RS were collected on 0, 2, 3, 4, 5, 6, 7, 9, 11, 14 and 20 pi, while serum was collected on days 0, 7, 14, and 20 pi. Two inoculated adult deer were euthanized on days 2 and 5 pi and the remaining animals were euthanized on day 20 pi. Following necropsy, multiple specimens including a tracheal swab, bronchial swabs, and several tissues (palatine tonsil, pharyngeal tonsil, nasal turbinate, medial retropharyngeal lymph node, mandibular lymph node, cerebellum, cerebrum, olfactory lobes, caudate nucleus, parotid salivary gland, thymus, trachea, bronchus, lung [4 sites], heart, tracheobronchial lymph node, mediastinal lymph node, liver, spleen, kidney, ileum, ileocecal valve, spiral colon, and mesenteric lymph node) were collected. Samples were individually bagged, placed on dry ice, and transferred to a −80 °C freezer until testing. Additionally, tissue samples were collected and processed for standard microscopic examination, a subset were also processed by *in situ* hybridization (ISH) and immunohistochemistry (IHC). For this, tissue sections of approximately ≤0.5 cm in width were fixed by immersion in 10% neutral buffered formalin (≥20 volumes fixative to 1 volume tissue) for approximately 24 h, and then transferred to 70% ethanol, followed by standard paraffin embedding techniques. Slides for standard microscopic examination were stained with hematoxylin and eosin (HE).

### Nucleic acid extraction and real-time RT-PCR (rRT-PCR)

Nucleic acid was extracted from nasal and oral secretions, feces, and all the tissue samples collected at necropsy. Before extraction, 0.5 g of tissues were minced with a sterile scalpel and resuspended in 5 ml DMEM (10% w/v) and homogenized using a stomacher (Stomacher^®^ 80 Biomaster; one speed cycle of 60s). Homogenized tissue samples were cleared by centrifugation (2000 × *g* for 10 min) and 200 µL of the homogenate supernatant used for RNA extraction using the MagMax Core extraction kit (Thermo Fisher, Waltham, MA, USA) and the automated KingFisher Flex nucleic acid extractor (Thermo Fisher) following the manufacturer’s recommendations. The real-time reverse transcriptase PCR (rRT-PCR) was performed using the EZ-SARS-CoV-2 Real-Time RT-PCR assay (Tetracore Inc., Rockville, MD, USA), which detects both genomic and subgenomic viral RNA targeting the virus nucleocapsid protein (N) gene. The limit of detection (LOD) for this assay was previously establish as 250 viral genome copies per mL, and 1.75 viral genome copies per reaction [39]. An internal inhibition control was included in all reactions. Positive and negative amplification controls were run side-by-side with test samples.

### Virus Isolation and titrations

Nasal and oral secretions, feces, and tissues collected during the necropsy that tested positive for SARS-CoV-2 by rRT-PCR were subjected to virus isolation under biosafety level 3 conditions at Cornell University. Twenty-four well plates were seeded with ~75,000 Vero E6/TMPRSS2 cells per well 24 h prior to sample inoculation. Cells were rinsed with phosphate buffered saline (PBS) (Corning^®^) and inoculated with 150 µl of each sample and inoculum adsorbed for 1 h at 37 °C with 5% CO_2_. Mock-inoculated cells were used as negative controls. After adsorption, replacement cell culture media supplemented as described above was added, and cells were incubated at 37 °C with 5% CO_2_ and monitored daily for cytopathic effect (CPE) for 3 days. SARS-CoV-2 infection in CPE-positive cultures was confirmed with an immunofluorescence assay (IFA) as described previously [10]. Cell cultures with no CPE were frozen, thawed, and subjected to two additional blind passages/inoculations in Vero E6/TMPRSS2 cell cultures. At the end of the third passage, the cells cultures were subjected to IFA. Positive samples on viral isolation were subjected to end point titrations by limiting dilution using the Vero E6/TMPRSS2 cells. Forty-eight hours later, the titration plates were fixed and subjected to the IFA to determine the virus titers using the Spearman and Karber’s method and expressed as TCID_50_.ml^−1^.

### In situ hybridization (ISH)

Paraffin-embedded tissues were sectioned at 5 µm and subjected to ISH using the RNAscope ZZ probe technology (Advanced Cell Diagnostics, Newark, CA). *In situ* hybridization was performed to detect tissue distribution of SARS-CoV-2 RNA in tissues from tissue tropism study (tissue distribution and tropism of SARS-CoV-2 in WTD). Nasal turbinate, palatine tonsil, medial retropharyngeal, mandibular and tracheobronchial lymph nodes, and lung were tested by RNAscope 2.5 HD Reagents–RED kit (Advanced Cell Diagnostics) as previously described [10]. Proprietary ZZ probes targeting SARS-CoV-2 RNA (V-nCoV2019-S probe ref # 8485561) or anti-genomic RNA (V-nCoV2019-S-sense ref # 845701) designed and manufactured by Advance Cell Diagnostics were used for detection of viral RNA. A positive control probe targeted the *Bos taurus*–specific cyclophilin B (PPIB Cat# 3194510) or ubiquitin (UBC Cat # 464851) housekeeping genes, while a probe targeting dapB of *Bacillus subtilis* (Cat # 312038) was used as a negative control.

### Immunohistochemistry (IHC)

Paraffin-embedded tissues were sectioned at 5 µm and subjected to IHC using Vectastain Elite ABC Peroxidase (HRP) Kit (Vector Laboratories Cat # PK-6102) as previously described [40]. Tissues from study 2 (distribution and tropism of SARS-CoV-2 in WTD) including nasal turbinate, palatine tonsil, mandibular lymph nodes, medial retropharyngeal lymph nodes, trachea, lung, olfactory lobe, caudate nucleus of brain, and cerebellum, were subjected to IHC. Formalin-fixed paraffin-embedded (FFPE) tissues were deparaffinized with xylene and rehydrated through a series of graded alcohol solutions. Antigen unmasking was performed using Tris-based antigen unmasking solution (pH 9.0) by boiling the slides in the unmasking solution for 20 min (Vector Laboratories). Quenching of endogenous peroxidase was performed using 0.3% hydrogen peroxide solution (Abcam Cat # ab64218). A mouse monoclonal antibody targeting nucleoprotein (N) of SARS-CoV-2 (developed in Dr. Diel’s laboratory) was used as a primary antibody (SARS-CoV-2 NP mAb clone B61G11). For SARS-CoV-2 detection, tissue sections were incubated with anti-mouse biotinylated secondary antibody followed by incubation with the Vectastain Elite ABC HRP reagent. Finally, tissues sections were incubated with Vector DAB peroxidase substrate (Vector Laboratories Cat # SK-4100). Counterstaining was performed using Hematoxylin QS Counter stain solution (Vector Laboratories Cat # H-3404).

### Serology

Neutralizing antibody responses to SARS-CoV-2 was assessed by a virus neutralization assay (VNA) performed under BSL-3 conditions at Cornell. Twofold serial dilutions (1:8 to 1:1,024) of serum samples were incubated with 100 - 200 TCID_50_ of SARS-CoV-2 isolate NYI67-20 for 1 h at 37 °C. Following incubation of serum and virus, 50 µl of a cell suspension of Vero cells was added to each well of a 96-well plate and incubated for 48 h at 37 °C with 5% CO_2_. The cells were fixed and permeabilized as described above and subjected to IFA using a mouse mAb specific for the SARS-CoV-2 nucleoprotein (N) followed by incubation with a goat anti-mouse IgG (goat anti-mouse IgG, DyLight^®^594 Conjugate, Immunoreagent Inc.). Unbound antibodies were washed from cell cultures by rinsing the cells PBS, and virus infectivity was assessed under a fluorescence microscope. Neutralizing antibody titers were expressed as the reciprocal of the highest dilution of serum that completely inhibited SARS-CoV-2 infection/replication. Normal serum (from uninfected WTD) and convalescent WTD serum from a previous study [10] were used as a negative and positive controls, respectively.

### SARS-CoV-2 MinION whole genome sequencing (WGS) and genetic analysis

The genetic make-up of SARS-CoV-2 following replication in WTD over viral transmission was investigated by whole genome sequencing. For this, nasal secretions collected on days 2 to 9 pi from inoculated and oronasal secretions on days 2 and 3 pc in contact animals were subjected to MinION-based targeted SARS-CoV-2 WGS. A multiplex RT-PCR was developed as previously described [41] following the amplicon-based approach used by the ARTIC Network for sequencing SARS-CoV-2 (https://artic.network/ncov-2019). The primers used here target approximately 1,500-bp and 500-bp products with 100 bp of overlap between different amplicons, covering the entire SARS-CoV-2 genome. Primer sequences are available at <u>dx.doi.org/10.17504/protocols.io.brkxm4xn</u>, and the first-strand synthesis and PCR conditions are available at <u>dx.doi.org/10.17504/protocols.io.br54m88w</u>. Libraries were generated using the Native Barcode Kit, EXP-NBD104, Ligation Sequencing Kit, SQK-SQK109 (Oxford Nanopore Technologies [ONT]) and sequenced on a R9.4 flow cell for 6 hours. Raw reads were base called and demultiplexed with the MinIT device (ONT) and then processed through the artic-ncov2019-medaka conda environment (https://github.com/artic-network/artic-ncov2019) to obtain final consensus sequences and reads alignment files. Low frequency variants were initially called using LoFreq [42] and subsequently filtered using Variabel [43].

### ACE2 and TMPRSS2 transcription and expression

To assess the ACE2 and TMPRSS2 transcription, RNA samples extracted from nasal turbinate, palatine tonsil, and lungs from all 15 fawns from the transmission dynamics study (euthanized on days 21 or 22) and from 4 deer from the pathogenesis study (euthanized on day 20) were included. were treated with DNA-free^TM^ Kit DNase Treatment & Removal (Invitrogen) according the manufacturer’s instructions. The amount of RNA in each sample was measured using Qubit^TM^ RNA BR Assay Kit (Invitrogen) and then all the samples of nasal turbinate/palatine tonsil and lungs were diluted in ddH_2_O in order to obtain a concentration of 40 ng/ul and 10 ng/ul, respectively. Standard curves were prepared from a pool of RNA of all samples (2-fold dilutions). Custom primers and probe were designed to angiotensin-converting enzyme 2 (ACE2), transmembrane serine protease 2 (TMPRSS2) and glyceraldehyde 3-phosphate dehydrogenase (GAPDH) of WTD using PrimerQuest Tool from Integrated DNA Technologies website. The primers and probe sequence for WTD ACE2 were 5’-GGATCTTGGCGTACAGAGAAAG-3’, 5’-CTGTTCTTCAGTGGTGGATTGA-3’ and /56-FAM/TTCTGGCTC/ZEN/CTTCTCAGCCTTGTT/3IABkFQ/ based on *Odocoileus virginianus texanus* ACE2 accession number XM_020913306.1; for WTD TMPRSS2 were 5’-CCTGTATGTCTTGGCCCTTT-3’, 5’-CCTGGTCAGACGAAGGTTATG-3’ and /56-FAM/TGTGCAGAC/ZEN/CCTGTGGCTCTACTA/3IABkFQ/ based on *Odocoileus virginianus texanus* TMPRSS2 accession number XM_020907939.1; and for WTD GAPDH were 5’-TGAGATCAAGAAGGTGGTGAAG-3’, 5’-GCATCGAAGGTAGAAGAGTGAG-3’ and /56-FAM/CCAGGTTGT/ZEN/CTCCTGCGACTTCAA/3IABkFQ/ based on *Odocoileus virginianus texanus* glyceraldehyde-3-phosphate dehydrogenase (GAPDH) accession number XM_020878738.1. Real-time RT-PCR amplifications were performed in 20-μl reactions, with 5 μl of RNA, 10 μl of TaqMan^®^ RT-PCR mix (TaqMan® RNA-to-CtTM 1-Step Kit, Applied Biosystems), 0.5 μl of TaqMan^®^ RT-Enzyme Mix, 1 μl of the mixture of primers and probe (PrimeTime qPCR probe assays, Integrated DNA Technologies Inc.), and 3.5 μl of ddH_2_O. Amplification and detection were performed using CFX96 TouchTM Real-Time PCR Detection System (Bio-Rad) under following conditions: 15 min at 48 °C for reverse transcription, 10 min at 95 °C for polymerase activation and 40 cycles of 15 s at 95 °C for denaturation and 1 min at 60 °C for annealing and extension. The measurement of gene expression was performed by using the relative quantitation method (Wong, M. L., and J. F. Medrano. 2005. Real-time PCR for mRNA quantitation. Biotechniques39:75-85). Relative genome copy numbers were calculated based on the standard curve determined for each gene within CFX MaestroTM software (Bio-Rad), and expression levels of the genes tested were normalized to the housekeeping gene GAPDH. (The amount of relative mRNA detected in each sample was expressed as log2 genome copy number). Statistical analysis was performed by 2way ANOVA followed by multiple comparisons and by unpaired t-test. Statistical analysis and data plotting were performed using the GraphPad Prism software (Version 9.0.1).

To assess ACE2 and TMPRSS2 expression levels, nasal turbinate and palatine tonsil from controls (uninoculated) WTD were sectioned at 5 µm and subjected to IFA. Formalin-fixed paraffin-embedded (FFPE) tissues were deparaffinized with xylene and rehydrated through a series of graded alcohol solutions. Antigen unmasking was performed using Tris-based antigen unmasking solution pH 9.0 (Vector Laboratories ref # H-3301) by boiling the slides in the unmasking solution for 20 min. After 10 min at 0.2% Triton X-100 (in phosphate-buffered saline [PBS]) at room temperature (rt), and 30 min blocking using a goat normal serum (0.2% in PBS) at rt, tissues subjected to IFA. Then, were incubated for 45 min at rt using a rabbit polyclonal antibody (pAb) anti-ACE2 (Abcan ref # ab15348) and a mouse monoclonal antibody (mAb) anti-TMPRSS2 (Santa Cruz Biotechnology, Inc. ref # sc-515727). Followed by 30 min incubation at rt with a goat anti-rabbit IgG (goat anti-rabbit IgG, Alexa Fluor® 594) and a goat anti-mouse IgG antibody (goat anti-mouse IgG, Alexa Fluor® 488). Nuclear counterstain was performed with 4’,6-Diamidino-2-Phenylindole, Dihydrochloride (DAPI) (10 min at rt) and visualized under a confocal microscopy (LSM710 Confocal - Zeiss).

## ACKNOWLEDGMENTS

The authors thank the animal caretakers and veterinarians at NADC for their help and excellent care with the animals. We also thank the Animal Health Diagnostic Center at Cornell University for the use of extraction and real-time PCR equipment. Mention of tradenames or commercial products is solely for the purpose of providing specific information and does not imply recommendation or endorsement by the U.S. Department of Agriculture.

